# Hox genes limit germ cell formation in the short germ insect *Gryllus bimaculatus*

**DOI:** 10.1101/419119

**Authors:** Austen A. Barnett, Taro Nakamura, Cassandra G. Extavour

## Abstract

Hox genes are conserved transcription factor-encoding genes that specify the identity of body regions in bilaterally symmetrical animals. In the cricket *Gryllus bimaculatus*, a member of the hemimetabolous insect group Orthoptera, the induction of a subset of mesodermal cells to form the primordial germ cells (PGCs) is restricted to the second through the fourth abdominal segments (A2-A4). In numerous insect species, the Hox genes *Sex-combs reduced* (*Scr*), *Antennapedia* (*Antp*), *Ultrabithorax* (*Ubx*) and *abdominal-A* (*abd-A*) jointly regulate the identities of middle and posterior body segments, suggesting that these genes may restrict PGC formation to specific abdominal segments in *G. bimaculatus*. Here we show that all of these Hox genes, either individually or in segment-specific combinations, restrict PGC formation. Our data provides evidence for a segmental Hox code used to regulate the placement of PGC formation, reminiscent of the segmental Hox codes used in other arthropod groups to establish other aspects of segmental identity. These data also provide, to our knowledge, the first evidence for this ancient group of genes in determining PGC placement within the context of axial patterning in any animal studied thus far.

## Introduction

The Hox genes are an ancient family of transcription factor-encoding genes that play a conserved role in specifying the body regions of bilaterally symmetrical animals during development (reviewed in 1). In arthropods, Hox genes act to specify the distinct identities of different body segments (reviewed in 2), with mutations in Hox genes usually resulting in switches of segmental type called homeotic transformations (reviewed in 3). We previously showed that in the cricket *Gryllus bimaculatus*, which belongs to the hemimetabolous insect order Orthoptera, the primordial germ cells (PGCs) form from the mesoderm of the second to the fourth abdominal segments (A2-A4) (4) via a bone morphogenetic protein (BMP)-dependent mechanism (5). Given that BMP activity is not limited to the segments where PGCs form, but rather, is present in the dorsal regions of all body segments (5), these data suggested that some unidentified factor or factors must ensure that PGCs form specifically in A2-A4. As Hox genes play a conserved role in establishing segmental identity, here we test the hypothesis that Hox genes contribute to regulating the placement of the PGCs in A2-A4.

Data implicating Hox genes in embryonic germ line placement in other animal taxa are scarce. In the mouse *Mus musculus*, as in *G. bimaculatus*, PGCs are established via BMP signals that activate the transcription factor Blimp-1 (5-7). Mouse embryonic cells that take on PGC fate repress the Hox genes *Hoxa1* and *Hoxb1* via activity of the BMP-activated transcription factor Blimp-1 (7-9). This has been interpreted as reflecting the loss of somatic differentiation programs that is associated with adopting PGC fate. Repression of Hox genes during differentiation of human induced pluripotent stem cells (hIPS) into in vitro derived PGCs (iPGCs) (10) is consistent with the hypothesis that Hox gene expression and PGC fate are mutually exclusive. In a system where germ cells are specified by inductive signals from neighboring cells, the degree or robustness of the PGC differentiation response may be influenced by the degree of concomitant Hox gene knockdown. Indeed, it was recently reported report that among mouse embryonic cells expressing PGC markers, there are some cells that also express hematopoietic markers, including at least one Hox gene (11). One interpretation of these observations is that those putative PGCs that respond to inductive signals by expressing lower levels of germ cell markers, may still be prone to express some somatic markers. Thus, it may be that decreased levels of somatic markers like Hox genes can facilitate the acquisition of PGC fate.

A role for HoxD genes in patterning and elongation of the external genitalia in mice has long been recognized (12-15), and HoxA genes are required for the correct development and function of elements of the female reproductive system that are derived from the Müllerian ducts, including the uterus and endometrium (16-19). However, the functions of Hox genes known to be expressed in the genital ridges, the precursors to the gonads (see for example 20), has received less attention. Taken together, these data suggest that there may be an important, but poorly studied, role for Hox genes in the context of PGC establishment and gonad positioning in animals with inductive PGC specification.

In contrast to *G. bimaculatus*, in the model insect *Drosophila melanogaster* PGCs form early in development (21, 22), long before the activation of the Hox genes that establish the identities of the body segments (23-26). Following gastrulation, the PGCs migrate towards the somatic gonad precursors (reviewed in 27), which develop from the mesoderm of embryonic abdominal parasegments 10-12 (28). The Hox genes *abdominal-A* (*abd-A*) and *Abdominal-B* (*Abd-B*) are necessary for the formation of the gonad precursors, which is independent of PGC specification. Specifically, *abd-A* establishes anterior gonad fates, and *abd-A* and *Abd-B* act in concert to establish the posterior gonad fates (28-31). In addition, in adult male *D. melanogaster, Abd-B* is required for correct function of the accessory gland (32), which is a component of the reproductive system that regulates the female response to mating (33). Moreover, this Hox gene is also required to maintain the identity of both germ line and somatic stem cells in the adult testis (34-37). However, these somatic and post-embryonic functions of *Abd-B* do not affect embryonic PGC establishment in *D. melanogaster*, which takes places much earlier in development.

Across insects, the Hox genes *Sex-combs reduced (Scr), Antennapedia (Antp), Ultrabithorax (Ubx)* and *abdominal-A (abd-A)* have a conserved role in establishing the thoracic and abdominal segments during embryogenesis (reviewed in 2). To explore the role of these Hox genes in establishing *G. bimaculatus* PGCs, we used embryonic RNAi (eRNAi) to repress the function of each of these genes individually, and also in combination. We found that these Hox genes act within the abdomen in a segment-specific manner to restrict PGC formation. Reminiscent of their combinatorial action in specifying other aspects of segment identity, including ectodermal patterning and appendage differentiation, these data suggest that a “Hox code” is also needed for appropriate PGC specification. These are the first data, to our knowledge, demonstrating a role for these highly conserved and ancient genes in limiting germ line development in an animal, and provide evidence for an additional embryonic role of Hox genes outside of establishing anterior-posterior segmental identity.

### Results and Discussion

*G. bimaculatus* PGCs emerge from the lateral mesoderm of abdominal segments A2-A4, within the mesoderm of each hemisegment (*i.e.*, the left or right halves of the segment) bearing the PGCs (4). Four Hox genes, *Scr* (*Gb-Scr*), *Antp* (*Gb-Antp*), *Ubx* (*Gb-Ubx*) and *abd-A* (*Gb-abd-A*) (Figs. S1, S2), are expressed in the abdomen during *G. bimaculatus* embryogenesis (38, 39). To investigate whether these posterior Hox genes were expressed at a time and place that would enable them to supply segment-specific positional information for PGC development, we performed *in situ* hybridization during PGC formation. As previously described (38, 39), these Hox genes exhibit spatial collinearity in the ectoderm from gnathal to abdominal segments in wild type embryos at stage 8, a stage when all segments have been defined and are morphologically distinct (Fig. 1 and Fig. S3). Because PGCs arise from the abdominal mesoderm, we asked whether these Hox genes were also expressed in the mesoderm. *Gb-Scr, Gb-Antp* and *Gb-abd-A* transcripts were expressed in the mesodermal cells of the abdominal segments (Fig. 1B, C, E), whereas *Gb-Ubx* transcripts were not detected above background levels in the dorsal mesodermal cells of A2-A10, but rather appeared to be restricted to the ectoderm in these segments (Fig. 1D).

**Fig. 1.**
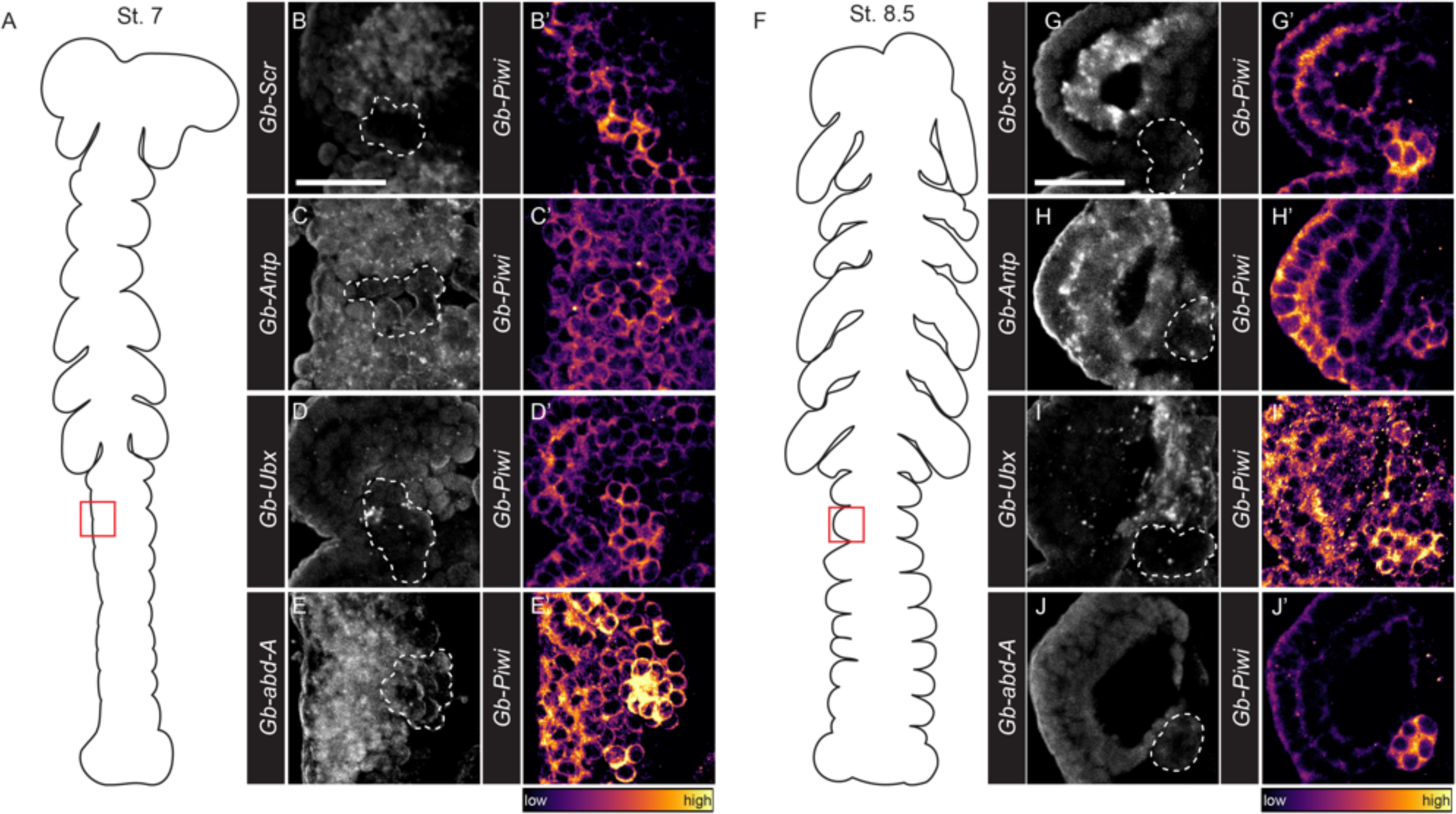
Posterior Hox gene expression adjacent to but absent from *G. bimaculatus* PGCs. (A,F) Schematic illustration of the embryonic stages (St.) (62) used for expression analysis. Red rectangle shows enlarged area in B-E and G-J’. *In situ* hybridization for *Gb-Scr, Gb*-*Antp, Gb*-*Ubx* and *Gb*-*abd-A* expression in the third abdominal segment (A3) at ES 7 (B-E) and ES 8.5 (G-J) of wild type embryos. (B’-J’) Co-staining with Gb-Piwi protein in A3. (B,G) Expression of *Gb-Scr* is enriched in the abdominal mesoderm and undetectable above background levels in the ectoderm. (B’,G’) Co-staining with Gb-Piwi protein reveals *Gb-Scr* expression is excluded from PGCs. (C,H) *Gb-Antp* expression is detected in abdominal mesoderm and ectoderm. (C’,H’) *Gb-Antp* transcript is undetectable in *G. bimaculatus* PGCs at St. 7 and at very low levels at St. 8.5. (D,I) Weak *Gb-Ubx* expression is observed in the ventral mesoderm of A3. (D’,I’) PGCs do not detectably express *Gb-Ubx*. (E,J) High *Gb-abd-A* expression is detected in the A3 ectoderm and mesoderm. (E’,J’) PGCs display lower expression levels of *Gb-abd-A* than the adjacent mesoderm. Scale bars = 100 μm in B (applies to B-E’), G (applies to G-J’).

To determine whether the mesodermally-expressed Hox genes were also expressed in PGCs, we conducted co-detection of Piwi protein, a PGC marker in *G. bimaculatus* (4), and Hox gene transcripts. We detected *Gb-Scr, Gb-Antp* and *Gb-abd-A* transcripts in mesodermal cells adjacent to PGCs at stages 7-8.5, but transcripts were undetectable or at very low levels in the PGCs themselves (Fig. 1B). Gb-Ubx expression was also undetectable in the PGCs at these stages (Fig. 1D, I). Taken together, these results suggest that, as in mice, *G. bimaculatus* Hox gene transcription is active in somatic cells adjacent to PGCs, but is repressed or inactive within PGCs (7-9).

Given their expression in cells in close proximity to PGCs, we hypothesized that these Hox genes might play a role in PGC formation, placement, or maintenance. To test this hypothesis, we used embryonic RNA interference (eRNAi) to abrogate the activity of one or more of these Hox genes during *G. bimaculatus* embryogenesis (Table S1), and assessed the effect on PGC number and segmental position. Specifically, we compared the PGC number per hemisegment and per embryo, as well as the presence or absence of PGC clusters in each hemisegment, to controls (Supplementary Materials and Methods; Tables S2-S7).

eRNAi against *Gb-Scr* efficiently depleted *Gb-Scr* transcripts as assessed by qPCR (Fig. S4), although this was not accompanied by transformation of gnathal appendages to a thoracic identity (n=36; Fig. 2L), as is often observed upon *Scr* knockdown in other insects (e.g. 40, 41). *Gb-Scr* eRNAi did, however, result in an increase in PGC number as well as a significant number of segments bearing PGC clusters in A2-A4 relative to controls (Fig. 2*A*-*B, F-G*). Furthermore, the total number of PGCs per embryo increased significantly relative to control injections (Fig. 2*K*).

**Fig. 2.**
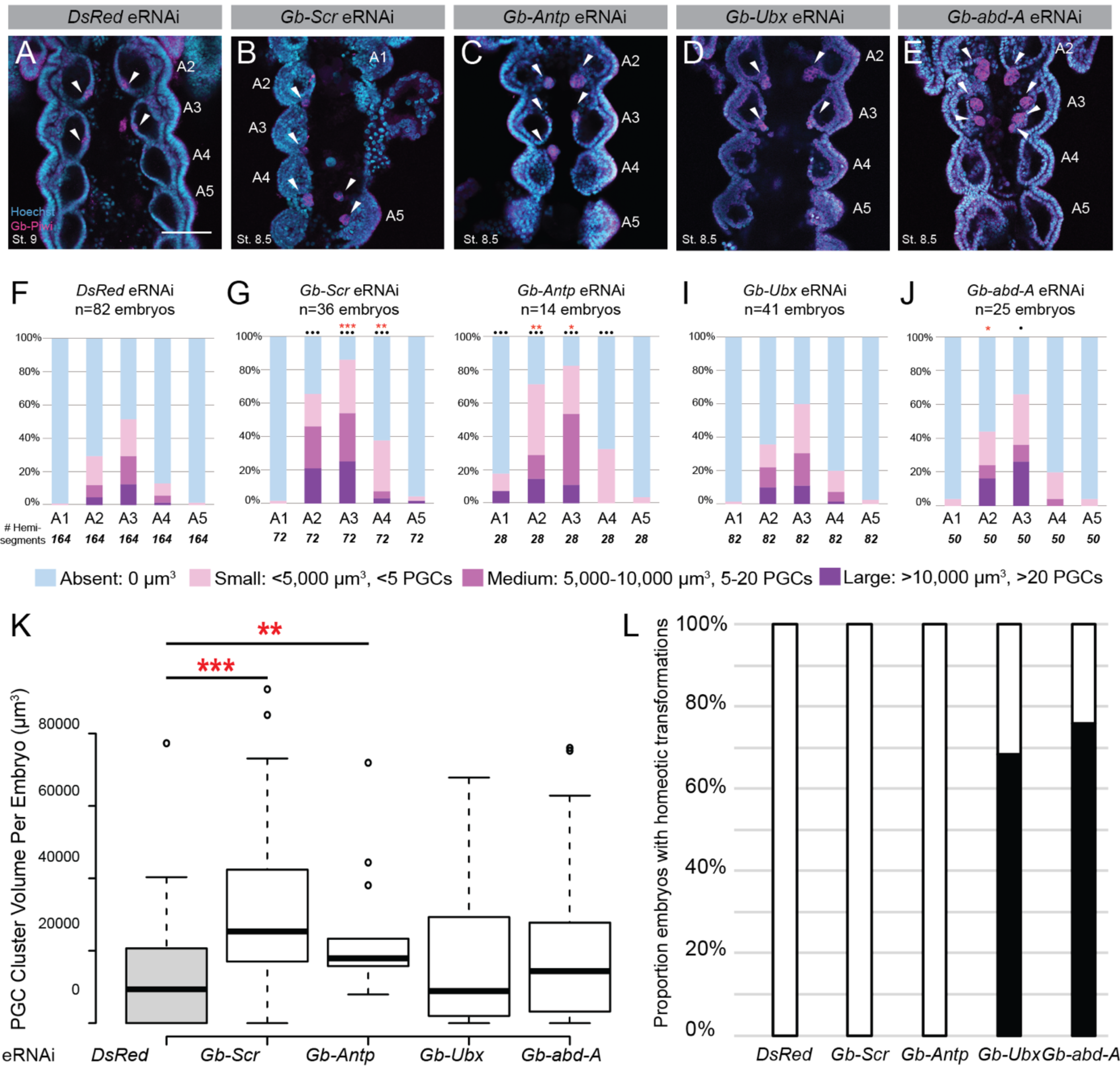
Embryonic RNA interference (eRNAi) of *Gb-Scr, Gb-Antp* and *Gb-abd-A* increases in PGC number. (*A-E*) Confocal images of the A2-A5 of a representative embryo from each knockdown condition as well as the control condition (*DsRed* eRNAi). Gb-Piwi (magenta) marks PGC clusters (arrowheads) (4). Anterior is up; scale bar = 100 μm. (*F-J*) PGC cluster quantifications of each eRNAi treatment (G-J) compared to (F) *DsRed* controls. Red asterisks denote significant size differences of PGC clusters in that segment compared to controls. Black dots denote significant differences in presence/absence of PGC clusters compared to controls. Numbers below each bar correspond to the number of hemisegments scored. (*K*) Box plot showing the distribution of total PGC volumes per embryo in each knockdown condition and the control condition (grey). Red asterisks denote significance levels resulting from a Mann-Whitney test. The whiskers extend to data points that are 1.5 times above the interquartile range away from the first or third quartile. The black lines represent the medians. n= the number of embryos observed for each condition. (*L*) 100% stacked bar chart showing the proportion of eRNAi embryos displaying homeotic phenotypes. Ax = abdominal segment x; St. = embryonic stage (62). All P-values for each test are listed in Tables *S2 and S4*. ^*/•^P<0.05, ^**/••^P<0.01, ^***/•••^P<0.001.

eRNAi against *Gb-Antp*, similarly to the *Gb-Scr* eRNAi treatment, did not result in a homeotic phenotype (n=14; Fig. 2L). However, our qPCR results showed that this treatment was also sufficient to reduce *Gb-Antp* transcripts (Fig. S4). *Gb-Antp* eRNAi significantly increased the proportion of segments bearing PGC clusters in A1-A4, and also increased PGC number in A2 (Figs. 2*C, H*) and overall (Fig. 2*K*). Thus, loss of *Gb-Antp* resulted both in additional PGCs in the correct segments, as did loss of *Gb-Scr*, and also in ectopic PGCs in A1.

*Gb-Ubx* eRNAi resulted in the transformation of A1 appendages (pleuropodia) towards ectopic walking legs (n=28/41; Figs. 2L, *SB5, S6B, F, S7A*), supported by expression of the appendage marker *Distal-less* (*Dll*) (42) (Fig. S6B) and reduced expression of the *G. bimaculatus* orthologue of *tramtrack* (*ttk*), a gene that shows enriched expression in pleuropodia (Fig. S6F), and consistent with *Ubx* knockdowns in other insects (43-46). Furthermore, qPCR showed that dsRNA injections reduced *Gb-Ubx* transcripts (Fig. S4). However, *Gb-Ubx* eRNAi did not significantly affect PGC number, or the numbers of PGC clusters, in any segment, or overall, relative to controls (Figs. 2*D, I* and *K*).

eRNAi targeting *Gb-abd-A* transcripts resulted in ectopic appendages throughout all abdominal segments (n=19/25; Figs. 2L, S*5C, S6C, G, S7B*). These ectopic outgrowths expressed *Dll* (Fig. S6C) but not *ttk* (Fig. S6G), and were consistent with outgrowths observed in *abd-A* knockdowns in other insects (43, 47-50). qPCR confirmed a decrease in *Gb-abd-A* transcripts following the eRNAi treatment (Fig. S4). *Gb-abd-A* eRNAi increased both PGC number in A2, and also the proportion of hemisegments bearing PGC clusters in A3 (Figs. 2*E, J* and *K*). Together, the results of these single Hox gene knockdowns suggest that *Gb-Scr* and *Gb-abd-A* represses mesodermal transformation to PGCs in A2-A3, and that *Gb-Antp* represses PGC formation in A1-A4.

In arthropods, Hox genes often work in concert to either activate or repress transcriptional targets (reviewed in 2). Therefore, we explored the possibility that the aforementioned Hox genes could be acting together in the context of PGC specification. We predicted that if a combination of Hox genes worked together to modulate PGC formation, a double knockdown of these genes would result in unique PGC defects relative to the defects observed in the single knockdowns discussed above. To test this prediction, we systematically injected embryos with equal amounts of dsRNA targeting each pairwise combination of these posterior *G. bimaculatus* Hox genes (Table S1).

Unexpectedly, all double eRNAi treatments that involved *Gb-Scr* dsRNA as a partner resulted in embryonic lethality one day after injection (Table S1). We therefore could not study its interaction with the other Hox genes eRNAi simultaneously targeting *Gb-Antp* and *Gb-Ubx* resulted in the same embryonic homeotic transformation as *Gb-Ubx* single knockdowns (*i.e.*, an ectopic fourth leg on A1, n=27/31; Fig. *S5D, S7C*). This double knockdown also resulted an increase in the presence of PGCs in A2-A5, as well as an increase in the number of PGCs in segments A2-A4 and overall compared to controls (Figs *3A, D, E*). Comparing these results to the *Gb-Ubx* and *Gb-Antp* single knock-downs, the overall PGC increase induced by the double knockdown (Fig. 3D) was not significantly different than that induced by *Gb-Antp* knockdown alone (Fig. 2K), suggesting that *Gb-Antp* act alone to repress PGC formation in A2-A4. However, the reduction in A1 PGC number in the double knockdown (Fig. 3A) relative to *Gb-Antp* alone (Fig. 2H), suggests that *Gb-Antp* acts to repress a PGC formation-promoting function of *Gb-Ubx* in A1. Furthermore, the presence of significantly more PGC clusters in A5 in the double knockdown (Fig. 3A, E) relative to both single knockdowns (Fig. 2H, I) also suggests that these genes act together to repress PGC formation in A5.

**Fig. 3.**
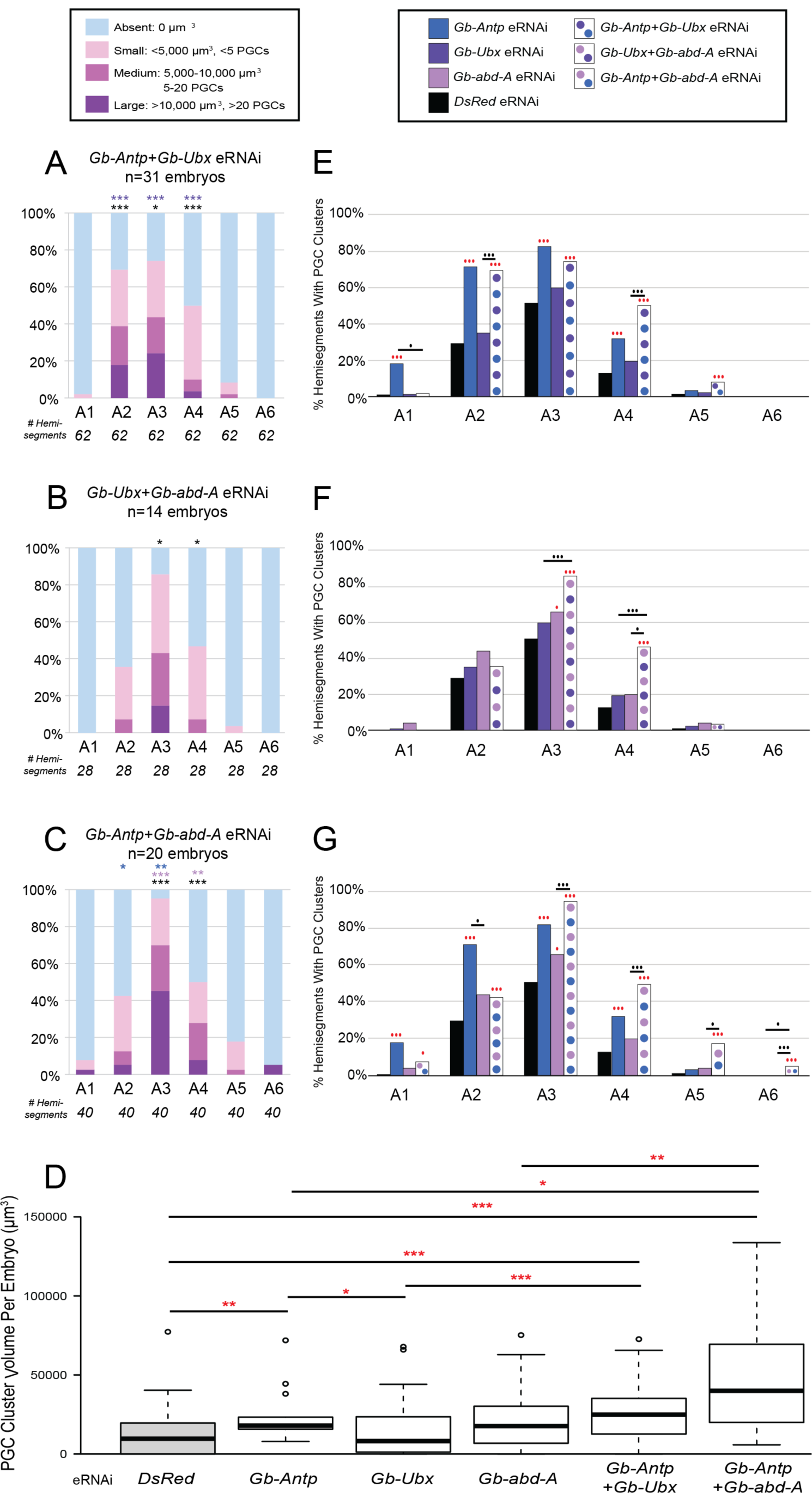
Double eRNAi reveals synergistic effects of Hox repression on germ cells. (*A-C*) PGC cluster quantifications of each eRNAi treatment. Asterisks denote significant size differences from *DsRed* controls (see Fig, 2F) and single eRNAi treatments (Fig. 2H-J) in that segment; asterisk colors indicate comparisons to *DsRed* (black), *Gb-Antp* eRNAi (dark blue), *Gb-abd-A* eRNAi (light purple), or *Gb-Ubx* eRNAi (dark purple). Numbers below each bar correspond to the number of hemisegments scored. (*D*) Box plot showing the distribution of total PGC volumes per embryo in each knockdown condition and the control condition (grey) except for *Gb-Ubx+Gb-abd-A* eRNAi, which displayed no significant total PGC differences in total (see text). Red asterisks denote significance levels resulting from a Mann-Whitney test. The whiskers extend to data points that are 1.5 times above the interquartile range away from the first or third quartile. The black lines represent the medians. n= the number of embryos observed for each condition. (*E-G*) Bar plots showing the proportion of hemisegments containing PGC clusters in single vs. double eRNAi treatments. The dots represent significance levels; red = compared to *DsRed* control; black = comparisons between single and double eRNAi conditions. All P-values for each test are listed in Tables *S3 and S5-S7*. ^*/•^P<0.05, ^**/••^P<0.01, ^***/•••^P<0.001.

RNAi simultaneously targeting *Gb-Ubx* and *Gb-abd-A* resulted in embryos with ectopic legs throughout the abdomen (n=8/14; Fig. *S5E, S7D*), exhibiting a similar phenotype to other studied insects (43, 49). These embryos also have significantly more PGC clusters in A3 and A4 compared to controls, as well as an increase in total PGC number in A4 (Figs. 3*B, F*). However, the overall increase in PGCs per embryo compared to controls was not statistically significant (Table S3). Comparing these results to the *Gb-Ubx* and *Gb-abd-A* single knockdowns suggests that *Gb-abd-A* alone is restricting PGC formation in A3, and that both *Gb-Ubx* and *Gb-abd-A* act together to restrict PGC formation in A4 (compare Fig. 2I, J with Fig 3*B*).

RNAi of *Gb-Antp* and *Gb-abd-A* together did not result in any detectable embryonic homeotic phenotype (data not shown). However, the effect on PGC formation in these embryos was striking: A1-A6 contained significantly more PGC clusters in A1-6 than controls (Figs. 3*C, D,* and *G*), and there were significantly more PGCs in A2-A4, and overall. Comparing this double knockdown to the single *Gb-Antp* and *Gb-abd-A* knockdowns revealed that the PGC increase caused by *Gb-Antp* eRNAi alone (Fig. 2H) was slightly suppressed in A2 (Fig. 3C), revealing a potential role of *Gb-Antp* in repressing *Gb-abd-A’s* ability to promote PGC formation in A2. Furthermore, this double knockdown provided evidence that *Gb-Antp* and *Gb-abd-A* act together to suppress PGC development in A3-A4, A6 and overall (Fig. *3D* and *G*).

Together, our results provide evidence for a Hox “code” specific to the formation of PGCs in *G. bimaculatus* abdominal segments (Fig. 4). We also propose that this “code” is at least partially separate from that encoding segmental identity, as the resulting Hox embryonic homeotic phenotypes do not always correlate with PGC positioning defects. For example, when we repress *Gb-abd-A* via eRNAi, A2-A3 bear pleuropodia-like appendages (Figs. *S5, S6*). In wild-type embryos, the pleuropodia are on A1, and thus we might expect that in this eRNAi condition, A2-A3 are transformed to an A1 identity. As the A1 segment generally lacks PGCs (4, 5), we should not observe PGCs in these segments in *Gb-abd-A* eRNAi injected embryos. However, we see an increase in PGCs in these segments in this condition (Fig. 2J). In another example, *Gb-Ubx+Gb-abd-A* eRNAi embryos bear ectopic leg-like structures on all abdominal segments (Fig. S5E). Being that *G. bimaculatus* wild-type embryos lack PGCs on their leg-bearing (thoracic) segments (4), we might expect an absence of Hox genes in A2-A4 in this eRNAi condition. However, we instead see an increase in the presence of PGCs in these segments (Fig. 3B).

**Fig. 4.**
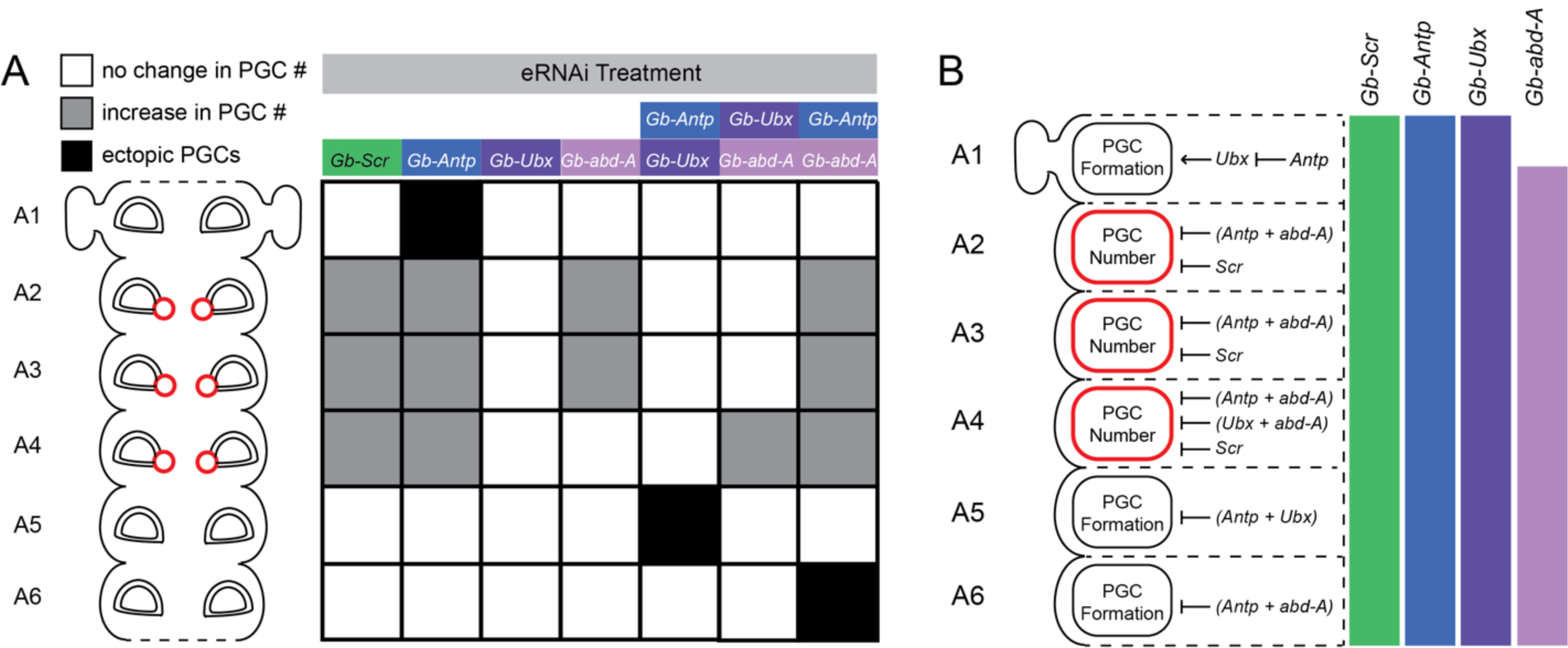
Combinatorial action of Hox genes regulate segmental positions of PGC. (*A*) Summary of the eRNAi results. In wild-type embryos, PGC clusters (red circles) form on A2-A4 (4). Significant changes in PGC number (grey) or appearance of ectopic PGCs (black) in each segment are induced by specific single or double Hox gene knockdowns. (*B*) Proposed mechanisms of combinatorial action of Hox genes on PGC development. Colored bars represent Hox expression domains (Fig. S3; (38, 39)). Arrows represent activating functions; and barred arrows represent repressive functions.

Hox codes that are used to pattern organs outside of the primary antero-posterior axes of animals have been described in other animal taxa. Examples include the use of Hox genes to pattern the proximo-distal axes of the tetrapod autopod (reviewed in 51), the branchial region of the vertebrate head (52), and the vertebrate neural tube (53). In insects, such non-axial Hox roles have also been previously described, such as those needed for the positioning of the bacteriomes in the seed bug *Nysius plebeius* (48), the formation of the lantern organ in the firefly *Photuris (sp.)* (54), and the differentiation of a specific accessory gland cell type in male fruit flies (32). Our results provide an example of Hox genes being used to coordinate cell type specification (PGCs) to ensure correct organ (gonad) positioning in the context of the body plan.

Our results suggest that Hox genes may play an indirect role in PGC specification. As has been suggested in the case of mouse PGC specification (7, 8, 55), downregulation of Hox genes may suppress somatic fate, permitting or facilitating adoption of PGC fate by BMP-responding cells. Alternatively, Hox genes may act in the somatic cells adjacent to PGCs to negatively regulate PGC fate, thus restricting the number of cells that can adopt PGC fate, or limiting the position where PGCs emerge in the abdominal segments.

The Hox code needed for the development of *G. bimaculatus* PGCs in A2-A4 may act to position these PGCs in an area where the gonad must form. *G. bimaculatus* PGCs do not undergo long-range migration (5), and the cells of the somatic gonad primordia are thought to lie within A2-A4 (4). To explore this possibility, we asked if *G. bimaculatus* orthologues of *Six4* and *eyes absent* (*eya*), gonad primordia markers used in *Drosophila* (56, 57), could identify putative somatic gonad precursors in *G. bimaculatus*. However, *in situ* hybridization did not show enhanced expression of either gene in cells adjacent to PGCs in A2-A4 (Fig. S8). These data suggest that either the gonad primordia are specified at later stages, or that these genes are not used to specify gonad primordia in *G. bimaculatus*. Further work will thus be needed to determine whether the increase in PGCs induced by many of the Hox eRNAi conditions is at the expense of mesoderm that would have been fated to be somatic gonadal cells in wild type embryos.

As in other animals with discrete gonads, arthropod PGCs must meet with somatic gonad cells and end up in a specific location in the body to form a functional gonad. In arthropods, the location of the gonad and often the location of PGCs when they first arise, is tied to specific body segments. In insects like *Drosophila*, where PGCs form much earlier than Hox gene activation and must migrate to the primordial gonad, the somatic gonad precursors rely on Hox genes to form in specific segments (28). In the beetle *Tenebrio molitor*, distinct PGC clusters form in many abdominal segments, and then coalesce into those segments that will ultimately contain the gonads (58). In the firebrat *Thermobia domestica* and the stick insect *Carausius morosus*, PGCs are thought to originate as a long cluster spanning multiple segments, before ultimately becoming confined to the gonads within specific segments (reviewed in 59). Outside of insects, in the spider *Parasteatoda tepidariorum*, the germ cells also arise as clusters, which are situated in the opisthosomal segments 2-6 (60). Taken together with the functional genetic data presented here, we suggest that assigning the PGC-bearing segments may be an ancestral role for Hox genes in arthropods.

## Materials and Methods

All Hox genes were cloned using a previously published *G. bimaculatus* transcriptome (61). The predicted translations of the resulting sequences were subjected to phylogenetic analysis to corroborate orthology (Fig. S2). Animal husbandry, eRNAi (Table S1), embryonic staging, cloning and qPCR (Table S8), *in situ* hybridizations and immunostainings, statistical tests and PGC quantifications were performed as previously described (4-6, 62). See Supplementary Material for detailed methods.

## Acknowledgments

We thank Dr. Taro Mito (University of Tokushima) for kindly allowing us access to draft genomic data for *G. bimaculatus*. This work was supported by NSF award IOS-1257217 to CGE.

## Supplementary Materials

### Supplementary Materials and Methods

#### Phylogenetic Analysis

Protein sequences of 119 annotated arthropod Hox gene (Fig. S1) and Distal-less (Dll) orthologues (for use as an outgroup) were retrieved from GenBank. These 119 sequences were used to make an alignment using MUSCLE (63) with eight iterations. The Smart Model Selection program (64) was used to find the best matrix for use in constructing the phylogeny (VT; AIC=164894.36226) and the best “decoration” (+G+I+F; AIC=164894.20896). This model was used with the resulting MUSCLE alignment to construct a maximum likelihood tree using PhyML (65). This resulted in a tree with a log likelihood of −81351.14064 (Fig. S2).

#### Quantitative PCR

Anterior abdominal segments A1-A5 were dissected from control or Hox gene eRNAi-treated embryos (n=5 per treatment for 2.5d and n=3 per treatment for 4d) using fine tungsten needles or fine forceps, and segments were pooled into single tubes. Total RNA was extracted using Trizol (Life Technologies) following the manufacturer’s directions. RNA pools were divided into two samples and each half was reverse transcribed to prepare cDNA using SuperScript III (Invitrogen). A no-reverse transcriptase control was performed in parallel for each sample. Each cDNA was divided into three replicate samples and used for qPCR. An MxP3005 machine (Stratagene) was used for qPCR as previously described (5). Relative transcript ratios in the qPCR study were calculated from experiments performed in triplicate and are shown as mean±s.d. in Fig. S4. The housekeeping gene *G. bimaculatus* β-*tubulin* was used as an internal control as previously described (5). Primers used are listed in Table S8.

## Supplementary Figure Legends

**Fig. S1.**
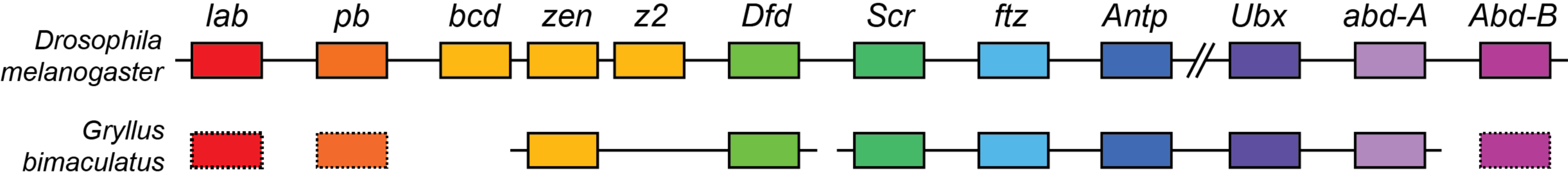
Putative chromosomal arrangement of the *G. bimaculatus* Hox complex. Schematic representation of the Hox complexes of *D. melanogaster* and *G. bimaculatus*. Orthologous genes are shown as color-coded boxes. Dashed rectangle represents genes found in *G. bimaculatus* transcriptome databases but not mapping to the *G. bimaculatus* draft genome database (Taro Mito, University of Tokushima, personal communication). Bar shows scaffold in *G. bimaculatus; gaps indicate regions where scaffold information is unavailable*. Double backslash in *D. melanogaster* cluster indicates the break between the Antennapedia complex and the Bithorax cluster complex. Illustration is not to scale.

**Fig. S2.**
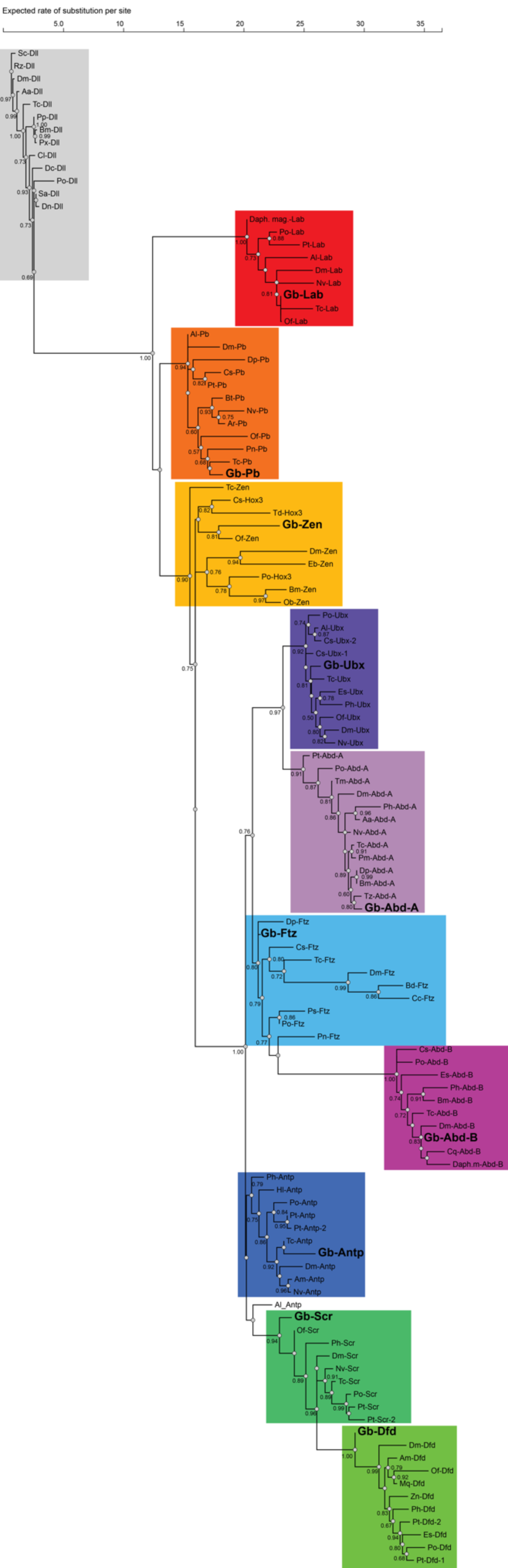
Maximum likelihood phylogeny of all *G. bimaculatus* predicted Hox amino acid sequences and other arthropod Hox amino acid sequences. Orthologous Hox clades are boxed and color-coded as in Figure S1. Distal-less (Dll) protein sequences were used as an outgroup. All nodes with approximate likelihood ratio tests (SH-like) support values with a value greater than or equal to 0.50 are labeled with the support value above the node. The tree resulted from an alignment of 119 arthropod amino acid protein sequences retrieved from GenBank. Protein abbreviations are as follows: Abd-A, Abdominal-A; Abd-B, Abdominal-B; Antp, Antennapedia; Dfd, Deformed; Ftz, Fushi tarazu; Pb, Proboscipedia; Scr, Sex combs reduced; Ubx, Ultrabithorax; Zen, Zerknullt. All prefixes to these Hox proteins denote species names, as follows: Aa, *Aedes albopictus*; Al, *Archegozetes longisetosus*; Am, *Apis mellifera*; Ar, *Athalia rosae*; Bd, *Bacrocera dorsalis*; Bm, *Bombyx mori*; Bt, *Bombus terrestris*; Cc, *Ceratitis capitata*; Cl, *Cimex lectularis*; Cq, *Culex quinquefasciatus*; Cs, *Cupiennius salei*; Daph. mag., *Daphnia magna*; Dc, *Diaphorini citri*; Dm, *Drosophila melanogaster*; Dn, *Diuraphis noxia*; Dp, *Daphnia pulex*; Eb, *Episyrphus balteatus*; Es, *Endeis spinosa*; Gb, *Gryllus bimaculatus*; Hl, *Habropoda laboriosa*; Mq, *Melipona quadrifasciata*; Nv, *Nasonia vitripennis*; Ob, *Operophtera brumata*; Of, *Oncopeltus fasciatus*; Ph, *Parhyale hawaiiensis*; Pm, *Papilio machaon*; Pn, *Paracyclopina nana*; Po, *Phalangium opilio*; Ps, *Pedetontus saltator*; Pt, *Parasteatoda tepidariorum*; Px, *Plutella xylostella*; Rz, *Rhagoletis zephyria*; Sc, *Stomoxys calcitrans*; Tc, *Tribolium castaneum*; Td, *Thermobia domestica*; Tm, *Tenebrio molitor*; Tz, *Trachymyrmex zeteki*; Zn, *Zootermopsis nevadensis*.

**Fig. S3.**
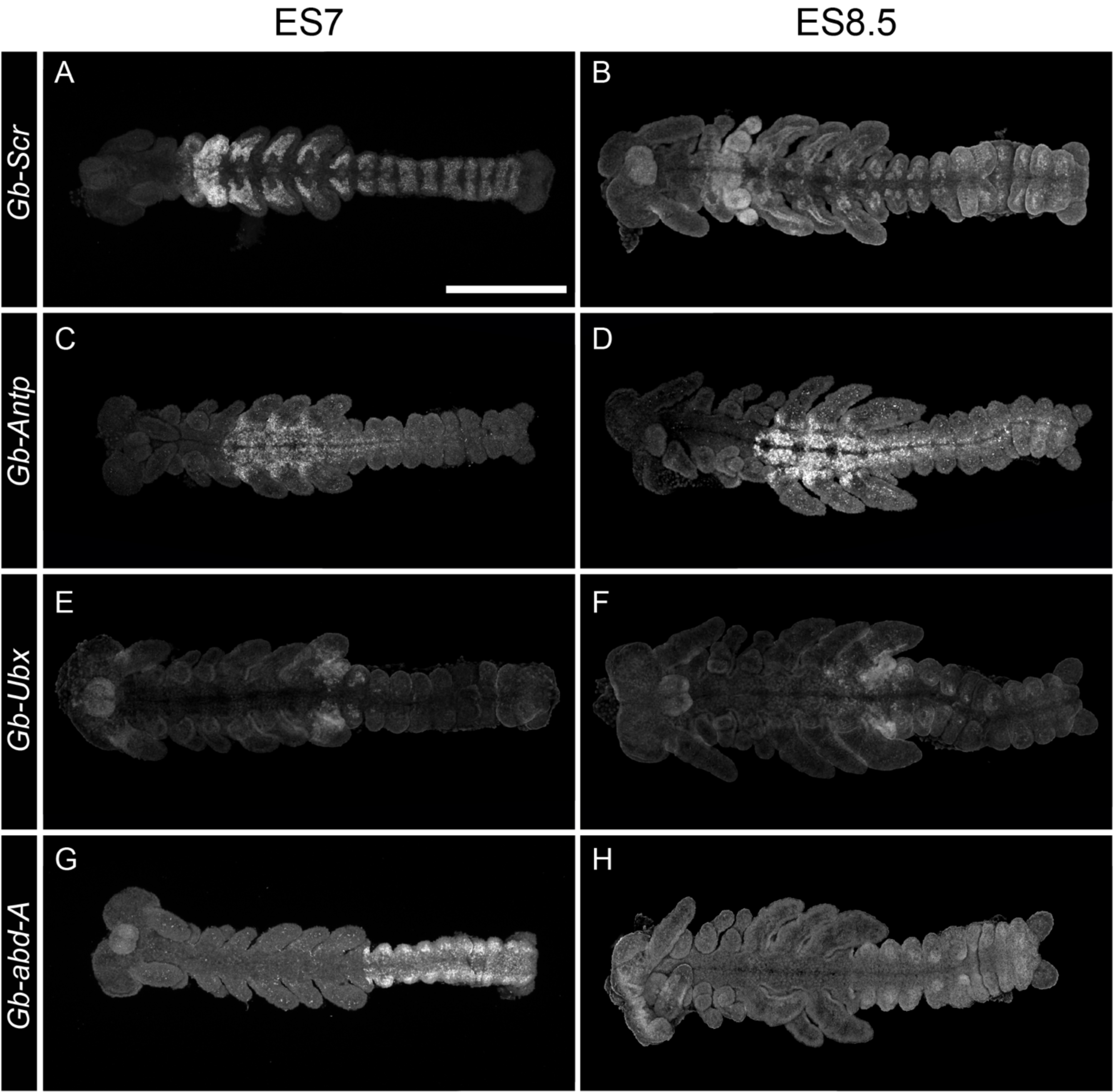
Expression of *Gb-Scr, Gb*-*Antp, Gb-Ubx* and *Gb-abd-A* transcripts during *G. bimaculatus* development. Fluorescent *in situ* hybridization of wild type embryos at egg stage (ES) ES7 (A, C, E, G) and ES8.5 (B, D, F, H). Egg staging as described in (62). Scale bar in A = 500 μm for all panels.

**Fig. S4.**
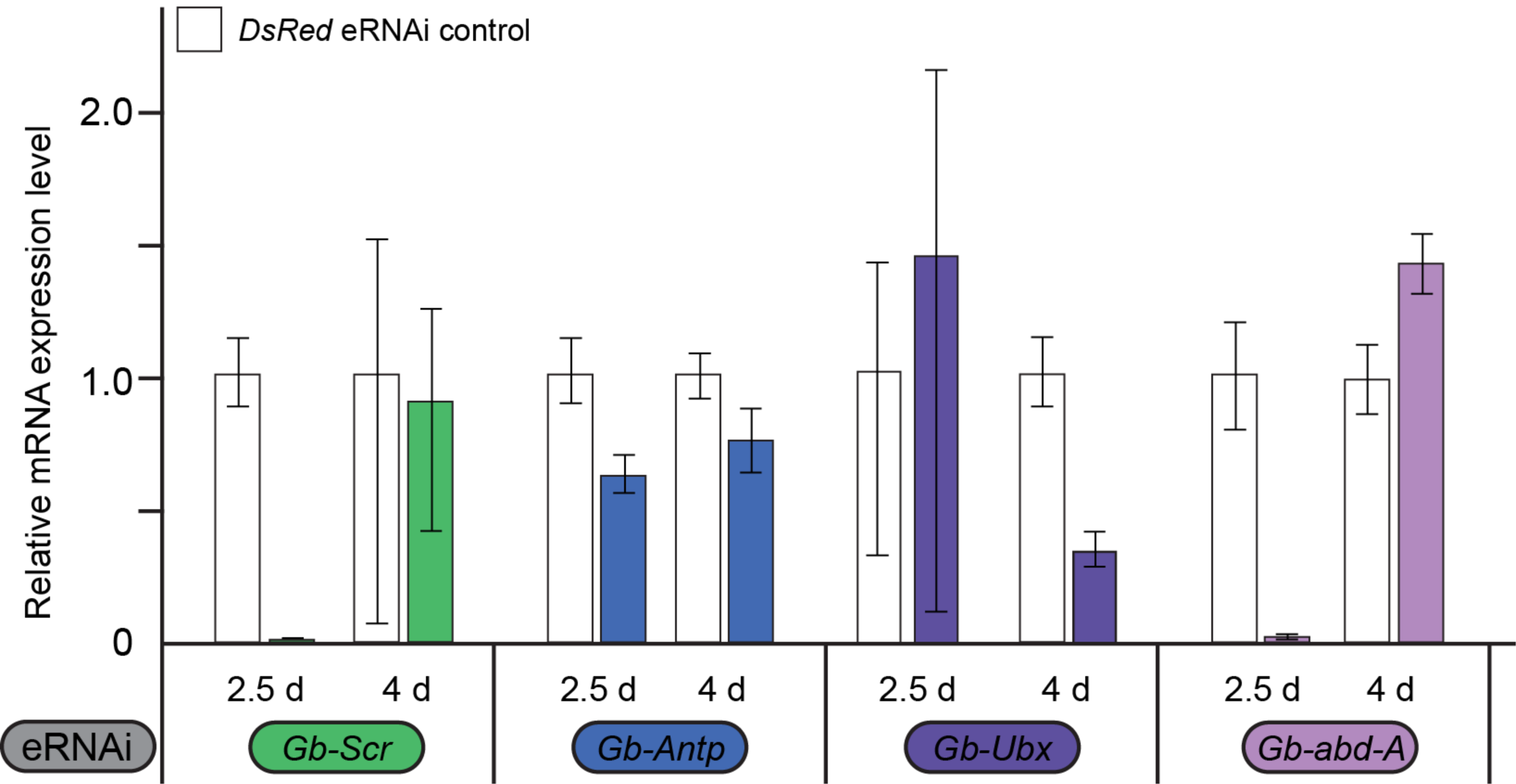
Validation of eRNAi knockdown via qPCR. Quantitative PCR was conducted to evaluate the efficacy of eRNAi-mediated knockdown against Hox genes in embryos at 2.5 days (d) and 4d after egg laying (AEL). At 2.5d AEL, PGC formation begins and at 4d AEL, PGC clusters are formed (4). Bars show relative expression levels normalized to *Gb-beta-tubulin*. eRNAi against all examined Hox genes reduced their transcript levels, if not at 2.5d AEL then by 4d AEL. Error bars show standard deviation.

**Fig. S5.**
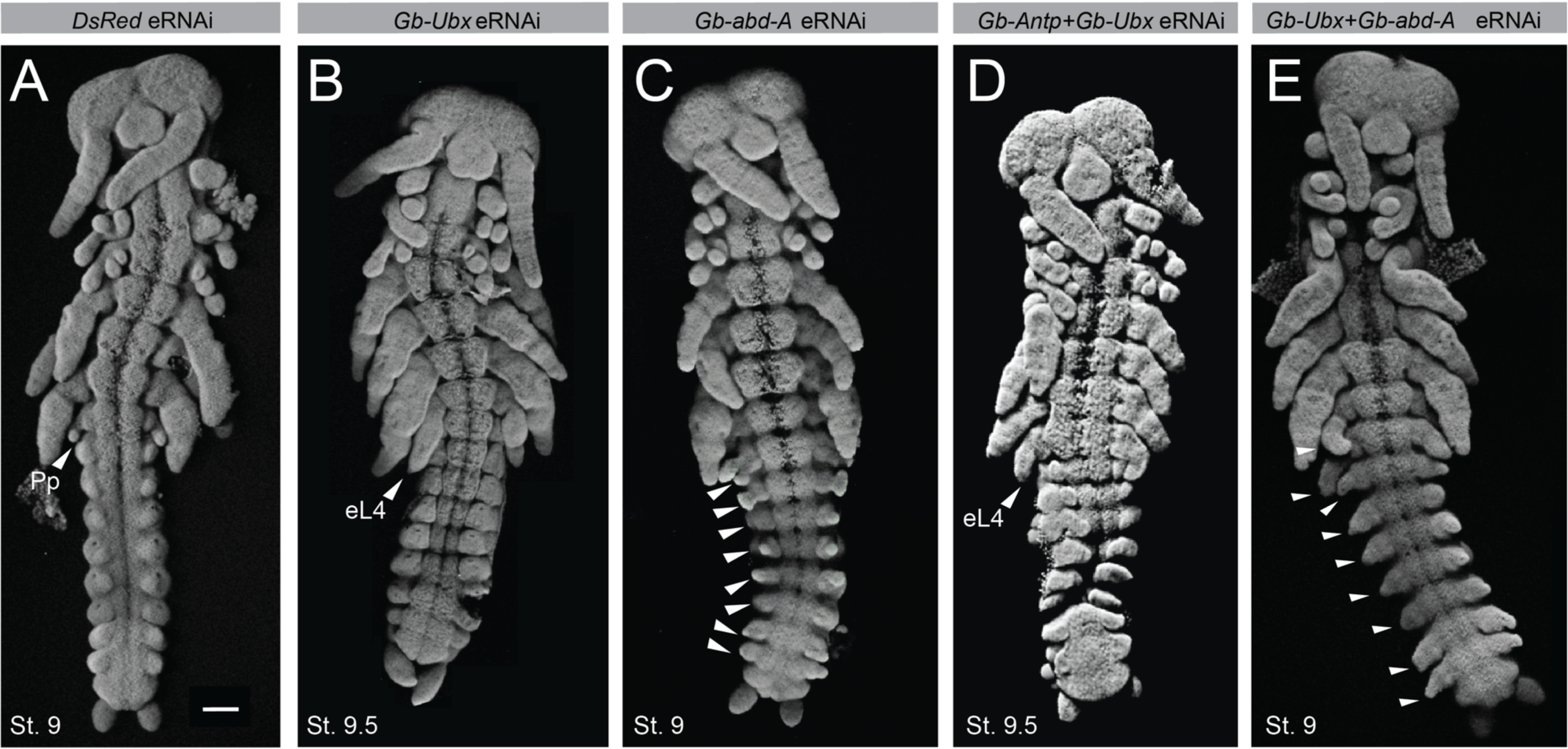
Homeotic transformations resulting from Hox eRNAi. (A) Example of an embryonic stage (St.) St.9 *DsRed* eRNAi control embryo showing the wild-type morphology, with pleuropodia (Pp) on the first abdominal segment (A1). (B) *Gb-Ubx* eRNAi resulted in the transformation of the pleuropodia into ectopic legs (eL4) on A1. (C) *Gb-abd-A* eRNAi resulted in the formation of ectopic pleuropodia-like appendages (arrowheads) on segments A2-A9. (D) Double eRNAi targeting *Gb-Antp* and *Gb*-*Ubx* resulted in the formation of ectopic legs (eL4) on A1. (E) Double eRNAi targeting *Gb-Ubx* and *Gb*-*abd-A* resulted in leg-like appendages (arrowheads) forming on segments A1-A9. Scale bars = 100 μm. All images are projections of optical confocal sections of embryos stained with Hoechst.

**Fig. S6.**
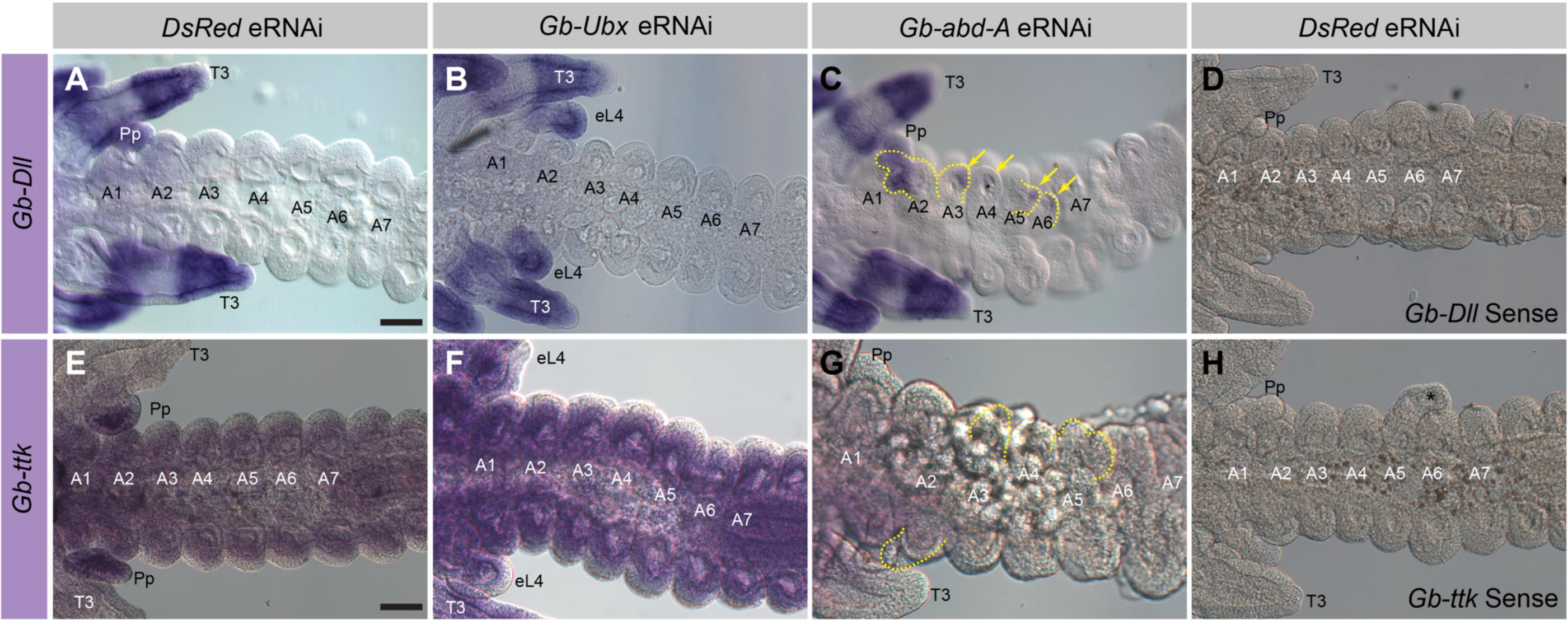
Expression patterns of *G. bimaculatus Distal-less* (*Gb-Dll*) and *tramtrack* (*Gb-ttk*) in *Gb-Ubx* and *Gb-abdominal-A* eRNAi embryos. (A-D) *In situ* hybridization for *Gb-Dll* in Hox eRNAi embryos compared with *DsRed* eRNAi control. (A) *Gb-Dll* expression in wild type embryos is detectable in appendages; thoracic legs and pleuropodia (A1) are visible in the figure. (B) *Gb-Ubx* eRNAi embryo shows enlargement of A1 appendage with distally localized *Gb-Dll* expression in the transformed structure. (C) *Gb-abd-A* eRNAi embryo shows ectopic extrusions (marked by yellow dotted lines) in A2-A6 segments. Ectopic *Gb-Dll* expression is detected in these extrusions. (D) Sense RNA probe against *Gb-Dll* used as negative *in situ* control in *DsRed* embryos. (E-H) *In situ* hybridization for *Gb-ttk* in Hox eRNAi embryos compared with *DsRed* eRNAi control. (E) *Gb-ttk* transcripts are enriched in pleuropodia, and low-level expression is also detectable throughout the entire embryo in *DsRed* eRNAi controls. (F) *Gb-Ubx* eRNAi embryo shows reduced expression in pleuropodia and slightly increased expression in the rest of the abdomen. (G) In *Gb-abd-A e*RNAi embryos *Gb-ttk* expression are lost from ectopic abdominal extrusions and nearly entirely from the rest of the embryo. (H) Sense RNA probe against *Gb-ttk* used as negative *in situ* control in *DsRed* embryos. Ax: abdominal segment x; eL4: ectopic 4^th^ leg; Pp: pleuropodia; T3: 3rd thoracic leg. Scale bar = 100 μm for all panels.

**Fig. S7.**
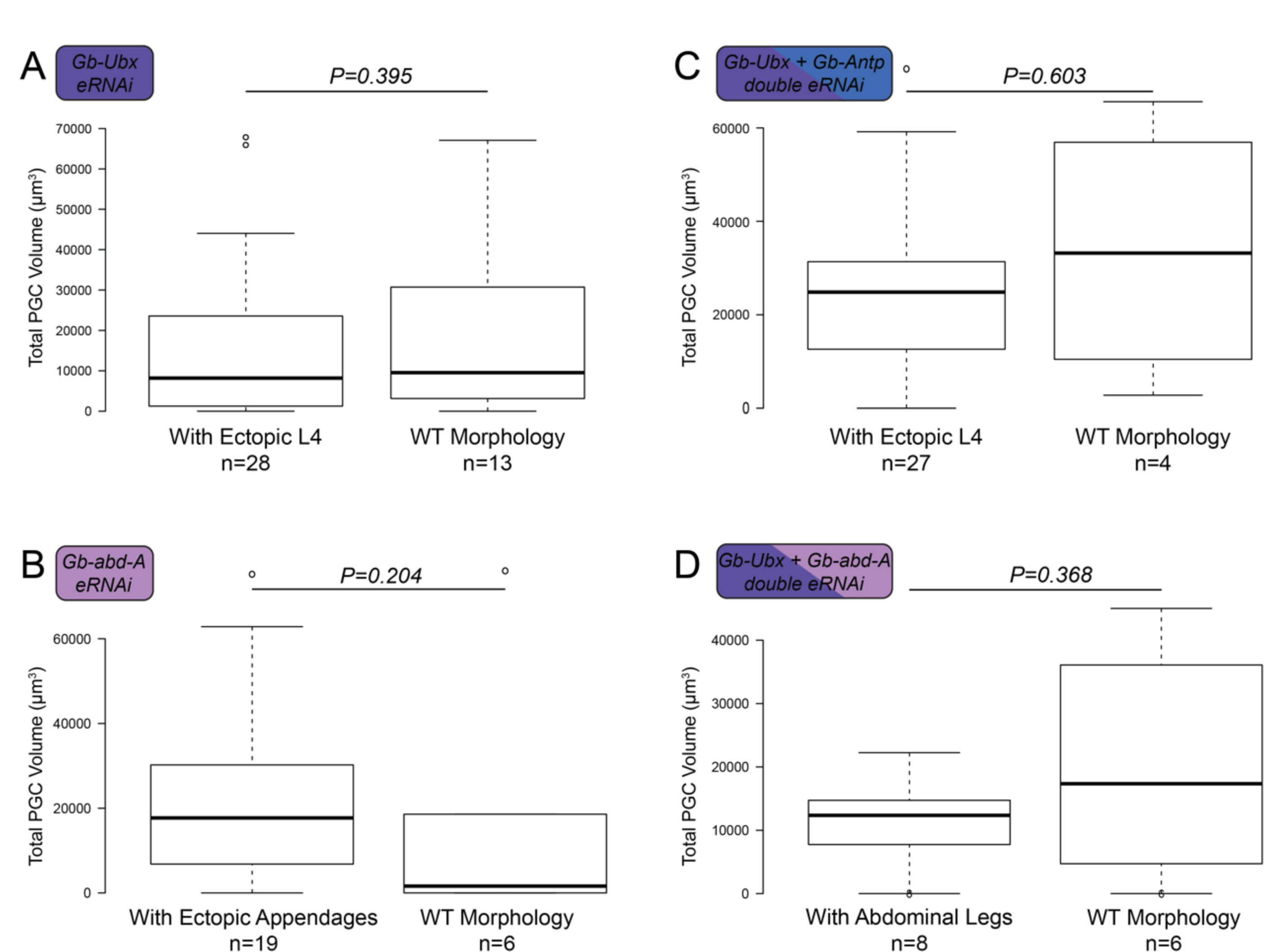
Statistical comparisons of PGC volumes from eRNAi embryos with and without homeotic phenotypes. (A) *Gb-Ubx*, (B) *Gb-abd-A* (C) *Gb-Antp+Gb-Ubx* and (D) *Gb-Ubx+Gb-abd-A* eRNAi knockdowns. In each condition, the distributions of the total PGC numbers per embryo were not statistically different between embryos with homeotic phenotypes and those without. The whiskers extend to data points that are 1.5 times above the interquartile range away from the first or third quartile. The black lines represent the medians. n= the number of embryos observed for each condition. P-values for (A-C) were based on the Mann-Whitney Test, and the P-value for (D) is based on a t-test due to the low number of samples in this comparison.

**Fig. S8.**
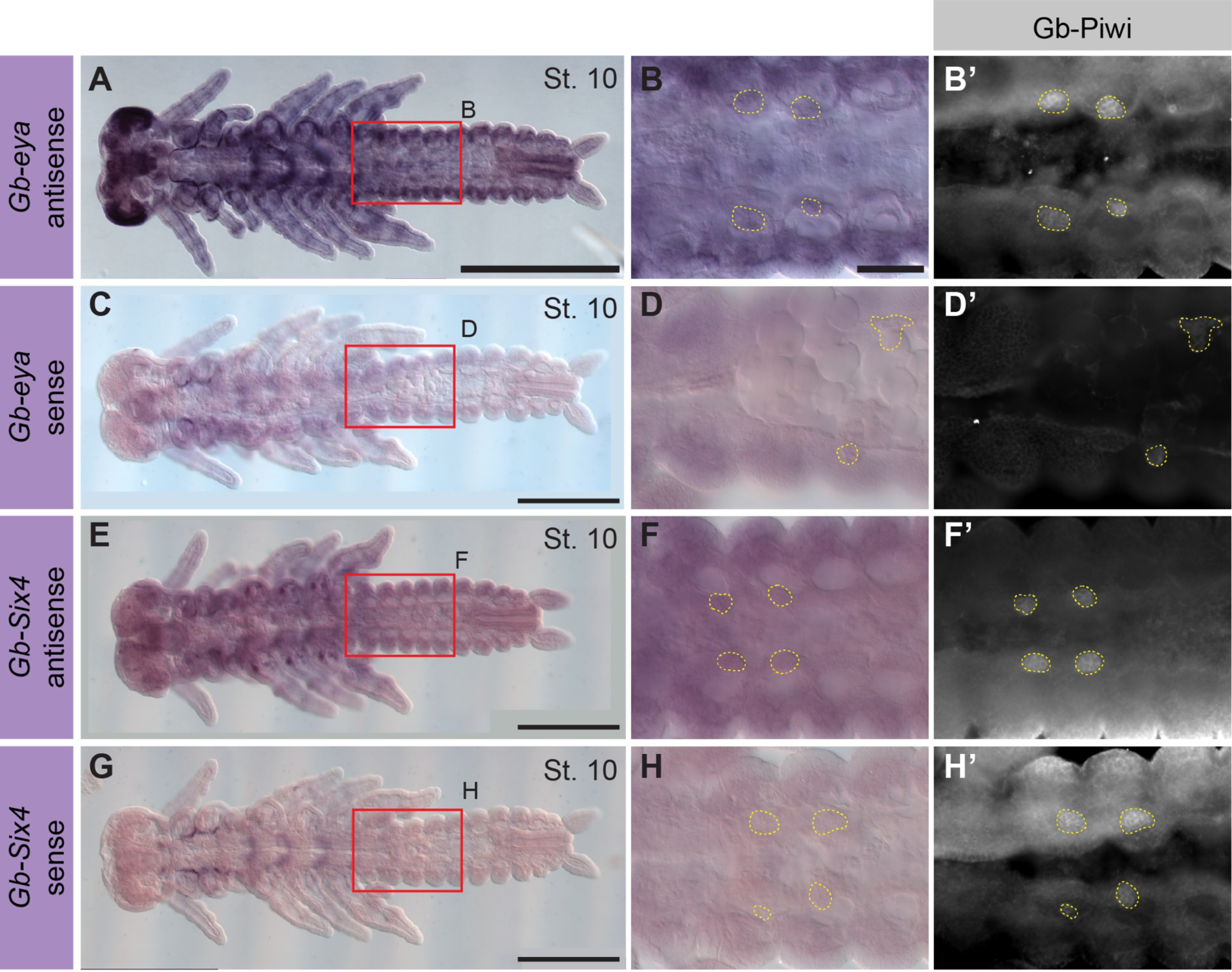
Expression of *G. bimaculatus eyes-absent* (*Gb-eya*) and *Six4* (*Gb-Six4*) in PGC development. (A-D’) *Gb-eya* transcripts are detectable at highest levels in the eye and brain primordia, and in the appendages in ring-like patterns. (B-B’) High magnification view of the abdominal region boxed in red in (A) shows that *Gb-eya* is expressed in each abdominal coelomic pouch, but not obviously at high levels in or adjacent to PGCs (marked by yellow dotted lines and labeled with anti-Gb-Piwi (B’)). (C, D) Sense RNA probe against *Gb-eya* was used as a negative *in situ* hybridization control. (E-H’) *Gb-Six4* transcripts are detected throughout the entire embryo, with increased expression in the thoracic coelomic pouches. (F-F’) High magnification view of the abdominal region boxed in red in (E) shows no enriched *Gb-Six-4* expression in or around PGCs. Scale bar = 500 μm in A, C, E, G and 100 μm in B, applicable to B-H’.

## Supplementary Tables

**Table S1:** Embryonic RNAi injection statistics.

| **<u>Injectant</u>** | **<u>Concentration</u>** | **<u># Embryos Injected</u>** | **<u># (%) Embryos Survived Injection to Stage 8</u>** | **<u># Embryos scored for PGCs in segments A1-A5</u>** |
| --- | --- | --- | --- | --- |
| ***DsRed dsRNA*** | 6 µg/µl | 3,190 | 1,395 (43.7) | 66 |
|  | 9 µg/µl | 417 | 176 (42.2) | 16 |
| ***Gb-Scr dsRNA*** | 6 µg/µl | 344 | 175 (50.9) | 36 |
| ***Gb-Antp dsRNA*** | 6 µg/µl | 1,103 | 251 (22.8) | 14 |
| ***Gb-Ubx dsRNA*** | 6 µg/µl | 203 | 163 (80.3) | 29 |
|  | 9 µg/µl | 372 | 112 (30.1) | 12 |
| ***Gb-abd-A dsRNA*** | 6 µg/µl | 1,126 | 375 (33.3) | 16 |
|  | 8 µg/µl | 149 | 22 (14.8) | 9 |
| ***Gb-Scr/abd-A RNAi*** | 3 µg/µl each | 268 | 0 (0) | 0 |
| ***Gb-Scr/Antp RNAi*** | 3 µg/µl each | 243 | 0 (0) | 0 |
| ***Gb-Antp/Ubx dsRNA*** | 3 µg/µl each | 374 | 201 (53.7) | 31 |
| ***Gb-Antp/abd-A dsRNA*** | 3 µg/µl each | 330 | 142 (43.0) | 20 |
| ***Gb-Ubx/abd-A dsRNA*** | 3 µg/µl each | 192 | 84 (43.8) | 14 |

**Table S2:**
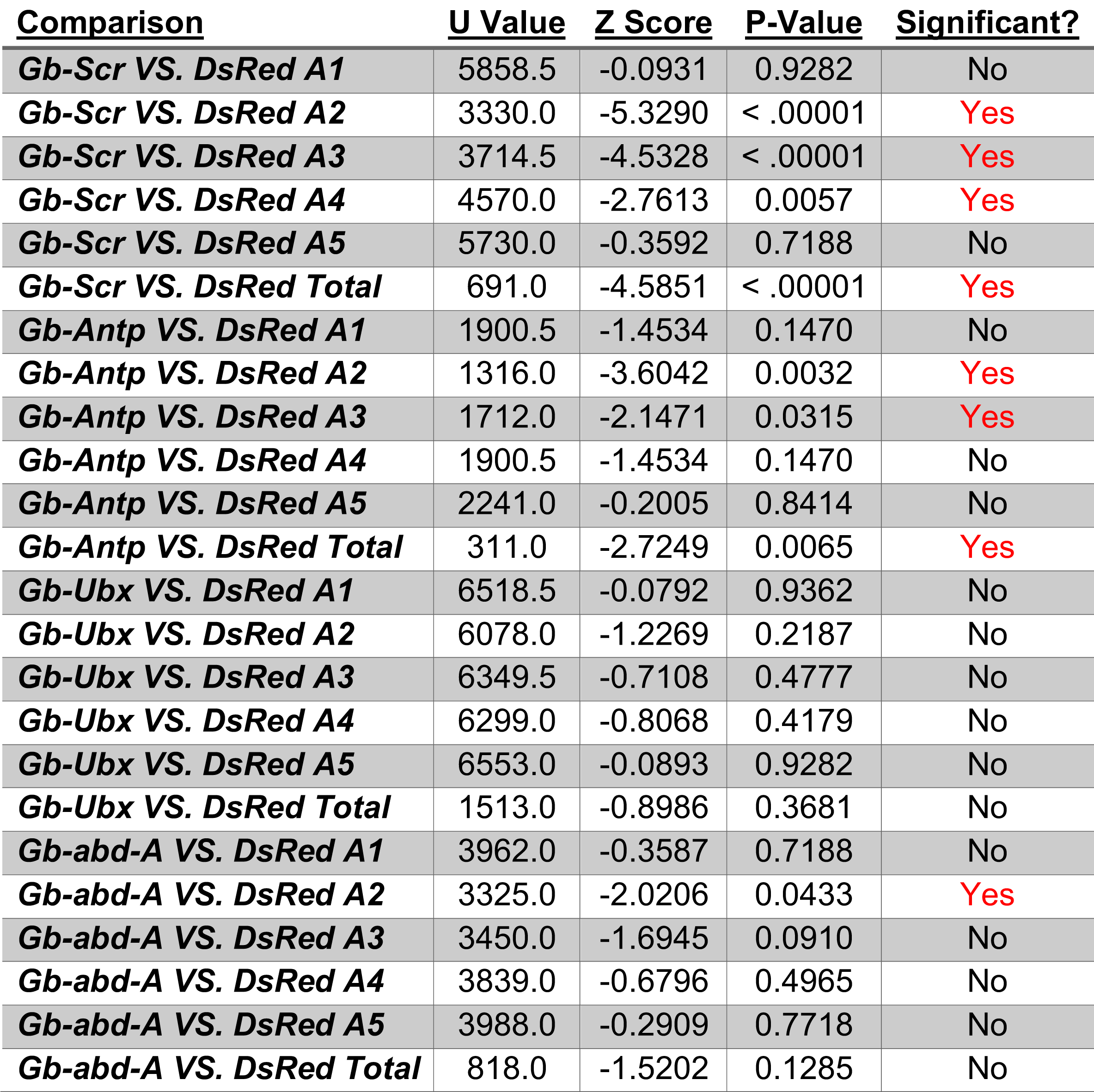
Mann-Whitney U test statistics on PGC measurements for single eRNAi treatments. A1-5, abdominal segments 1-5.

**Table S3:** Mann-Whitney U test statistics on PGC measurements for double eRNAi treatments. A1-5, abdominal segments 1-5.

| <u>Comparison</u> | <u>U Value</u> | <u>Z Score</u> | <u>P-Value</u> | <u>Significant?</u> |
| --- | --- | --- | --- | --- |
| *(Gb-Antp+Gb-Ubx) VS. DsRed A1* | 5033.5 | -0.1140 | 0.9124 | No |
| *(Gb-Antp+Gb-Ubx) VS. DsRed A2* | 2820.0 | -5.1610 | < .00001 | Yes |
| *(Gb-Antp+Gb-Ubx) VS. DsRed A3* | 3756.0 | -3.0268 | 0.0024 | Yes |
| *(Gb-Antp+ Gb-Ubx) VS. DsRed A4* | 3331.5 | -3.9947 | 0.0001 | Yes |
| *(Gb-Antp+ Gb-Ubx) VS. DsRed A5* | 4734.0 | -0.7969 | 0.4237 | No |
| *(Gb-Antp+ Gb-Ubx) VS. DsRed Total* | 660.0 | -3.9285 | 0.0001 | Yes |
| *(Gb-Ubx+ Gb-abd-A) VS. DsRed A1* | 2282.0 | 0.0496 | 0.9601 | No |
| *(Gb-Ubx+ Gb-abd-A) VS. DsRed A2* | 2204.0 | -0.3366 | 0.7279 | No |
| *(Gb-Ubx+ Gb-abd-A) VS. DsRed A3* | 1755.0 | -1.9888 | 0.0466 | Yes |
| *(Gb-Ubx+ Gb-abd-A) VS. DsRed A4* | 1598.5 | -2.5647 | 0.0105 | Yes |
| *(Gb-Ubx+ Gb-abd-A) VS. DsRed A5* | 2243.0 | -0.1931 | 0.8493 | No |
| *(Gb-Ubx+ Gb-abd-A) VS. DsRed Total* | 492.0 | -0.8460 | 0.3953 | No |
| *(Gb-Antp+ Gb-abd-A) VS. DsRed A1* | 3054.5 | -0.6721 | 0.5029 | No |
| *(Gb-Antp+ Gb-abd-A) VS. DsRed A2* | 2928.5 | -1.0485 | 0.2937 | No |
| *(Gb-Antp+ Gb-abd-A) VS. DsRed A3* | 1319.0 | -5.8563 | < .00001 | Yes |
| *(Gb-Antp+ Gb-abd-A) VS. DsRed A4* | 2040.0 | -3.7026 | 0.0002 | Yes |
| *(Gb-Antp+ Gb-abd-A) VS. DsRed A5* | 2744.0 | -1.5996 | 0.1096 | No |
| *(Gb-Antp+ Gb-abd-A) VS. DsRed A6* | 3116.0 | -0.4884 | 0.6241 | No |
| *(Gb-Antp+ Gb-abd-A) VS. DsRed Total* | 224.0 | -5.0191 | < .00001 | Yes |

**Table S4:**
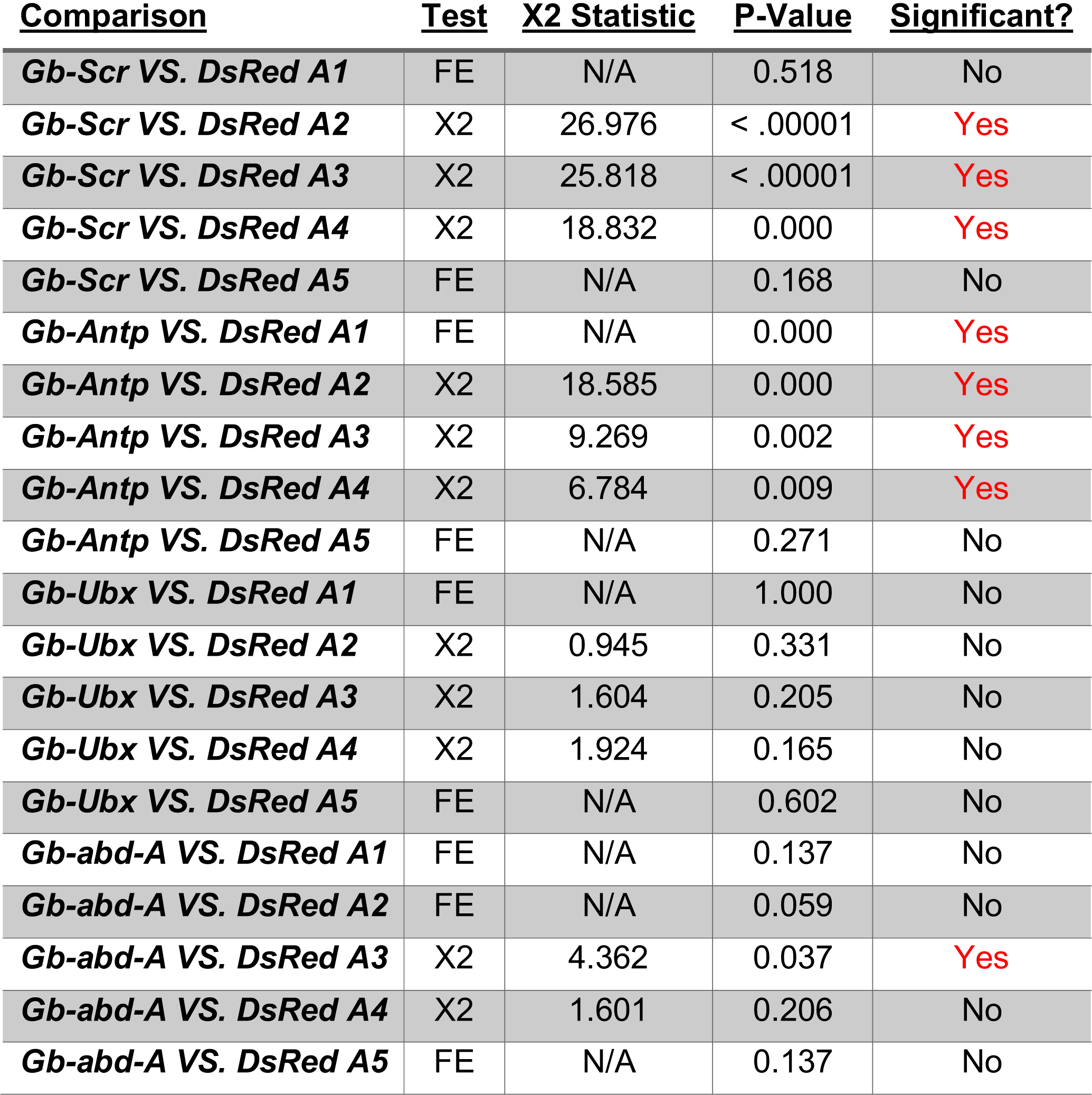
Statistical tests of presence/absence of PGC clusters in single Hox eRNAi treatments. FE = Fisher’s Exact Test; X2 = Chi-Square Test; N/A = not applicable; A1-5, abdominal segments 1-5.

**Table S5:** Statistical tests of presence/absence of PGC clusters in double Hox eRNAi treatments. FE = Fisher’s Exact Test; X2 = Chi-Square Test; N/A = not applicable; A1-6, abdominal segments 1-6.

| <u>Comparison</u> | <u>Test</u> | <u>X2 Statistic</u> | <u>P-Value</u> | <u>Significant?</u> |
| --- | --- | --- | --- | --- |
| *(Gb-Antp+ Gb-Ubx) VS. DsRed A1* | FE | N/A | 0.4743 | No |
| *(Gb-Antp+ Gb-Ubx) VS. DsRed A2* | X2 | 30.0584 | < .00001 | Yes |
| *(Gb-Antp+ Gb-Ubx) VS. DsRed A3* | X2 | 9.7186 | 0.0018 | Yes |
| *(Gb-Antp+ Gb-Ubx) VS. DsRed A4* | X2 | 35.1369 | < .00001 | Yes |
| *(Gb-Antp+ Gb-Ubx) VS. DsRed A5* | FE | N/A | 0.0065 | Yes |
| *(Gb-Ubx+ Gb-abd-A) VS. DsRed A1* | FE | N/A | 1.0000 | No |
| *(Gb-Ubx+ Gb-abd-A) VS. DsRed A2* | X2 | 0.4714 | 0.4923 | No |
| *(Gb-Ubx+ Gb-abd-A) VS. DsRed A3* | FE | N/A | 0.0007 | Yes |
| *(Gb-Ubx+ Gb-abd-A) VS. DsRed A4* | X2 | 18.5549 | 0.0000 | Yes |
| *(Gb-Ubx+ Gb-abd-A) VS. DsRed A5* | FE | N/A | 0.2710 | No |
| *(Gb-Antp+ Gb-abd-A) VS. DsRed A1* | FE | N/A | 0.0244 | Yes |
| *(Gb-Antp+ Gb-abd-A) VS. DsRed A2* | X2 | 11.2953 | 0.0008 | Yes |
| *(Gb-Antp+ Gb-abd-A) VS. DsRed A3* | X2 | 25.6402 | < .00001 | Yes |
| *(Gb-Antp+ Gb-abd-A) VS. DsRed A4* | X2 | 24.5565 | < .00001 | Yes |
| *(Gb-Antp+ Gb-abd-A) VS. DsRed A5* | FE | N/A | 0.0000 | Yes |
| *(Gb-Antp+ Gb-abd-A) VS. DsRed A6* | FE | N/A | 0.0000 | Yes |

**Table S6:** Mann-Whitney U test statistics on PGC measurements for single vs. double eRNAi treatments. A1-6, abdominal segments 1-6.

| <u>Comparison</u> | <u>U Value</u> | <u>Z Score</u> | <u>P-Value</u> | <u>Significant?</u> |
| --- | --- | --- | --- | --- |
| (Gb-Antp+ Gb-Ubx) VS. Gb-Antp A1 | 759.0 | -1.18816 | 0.234 | No |
| (Gb-Antp+ Gb-Ubx) VS. Gb-Antp A2 | 811.0 | 0.49243 | 0.624 | No |
| (Gb-Antp+ Gb-Ubx) VS. Gb-Antp A3 | 834.0 | 0.29197 | 0.772 | No |
| (Gb-Antp+ Gb-Ubx) VS. Gb-Antp A4 | 709.5 | 1.37706 | 0.168 | No |
| (Gb-Antp+ Gb-Ubx) VS. Gb-Antp A5 | 829.5 | 0.33119 | 0.741 | No |
| (Gb-Antp+ Gb-Ubx) VS. Gb-Antp Total | 194.0 | 0.55163 | 0.582 | No |
| (Gb-Antp+ Gb-Ubx) VS. Gb-Ubx A1 | 2470.5 | -0.03702 | 0.968 | No |
| (Gb-Antp+ Gb-Ubx) VS. Gb-Ubx A2 | 1764.5 | -3.13491 | 0.002 | Yes |
| (Gb-Antp+ Gb-Ubx) VS. Gb-Ubx A3 | 1997.5 | -2.19484 | 0.028 | Yes |
| (Gb-Antp+ Gb-Ubx) VS. Gb-Ubx A4 | 1824.5 | -2.89283 | 0.004 | Yes |
| (Gb-Antp+ Gb-Ubx) VS. Gb-Ubx A5 | 2339.0 | -0.57791 | 0.562 | No |
| (Gb-Antp+ Gb-Ubx) VS. Gb-Ubx Total | 399.0 | -2.68391 | 0.007 | Yes |
| (Gb-Ubx+ Gb-abd-A) VS. Gb-Ubx A1 | 1106.0 | 0.09464 | 0.928 | No |
| (Gb-Ubx+ Gb-abd-A) VS. Gb-Ubx A2 | 1063.0 | 0.57983 | 0.562 | No |
| (Gb-Ubx+ Gb-abd-A) VS. Gb-Ubx A3 | 944.0 | -1.39639 | 0.162 | No |
| (Gb-Ubx+ Gb-abd-A) VS. Gb-Ubx A4 | 864.0 | -1.94534 | 0.0511 | Approaching |
| (Gb-Ubx+ Gb-abd-A) VS. Gb-Ubx A5 | 1109.0 | -0.07361 | 0.944 | No |
| (Gb-Ubx+ Gb-abd-A) VS. Gb-Ubx Total | 258.0 | -0.55066 | 0.582 | No |
| (Gb-Ubx+ Gb-abd-A) VS. Gb-abd-A A1 | 672.0 | 0.28645 | 0.772 | No |
| (Gb-Ubx+ Gb-abd-A) VS. Gb-abd-A A2 | 586.0 | 1.18225 | 0.238 | No |
| (Gb-Ubx+ Gb-abd-A) VS. Gb-abd-A A3 | 658.5 | -0.42707 | 0.667 | No |
| (Gb-Ubx+ Gb-abd-A) VS. Gb-abd-A A4 | 531.0 | -1.75514 | 0.078 | No |
| (Gb-Ubx+ Gb-abd-A) VS. Gb-abd-A A5 | 697.0 | 0.02604 | 0.976 | No |
| (Gb-Ubx+abd-A) VS. Gb-abd-A Total | 163.0 | 0.33669 | 0.728 | No |
| (Gb-abd-A+ Gb-Antp) VS. Gb-abd-A A1 | 610.5 | 0.32864 | 0.741 | No |
| (Gb-abd-A+ Gb-Antp) VS. Gb-abd-A A2 | 891.0 | 0.88102 | 0.379 | No |
| (Gb-abd-A+ Gb-Antp) VS. Gb-abd-A A3 | 518.0 | -3.90977 | 0.000 | Yes |
| (Gb-abd-A+ Gb-Antp) VS. Gb-abd-A A4 | 661.0 | -2.74861 | 0.006 | Yes |
| (Gb-abd-A+ Gb-Antp) VS. Gb-abd-A A5 | 858.0 | -1.14898 | 0.250 | No |
| (Gb-abd-A+ Gb-Antp) VS. Gb-abd-A A6 | 950.0 | -0.40194 | 0.689 | No |
| (Gb-abd-A+ Gb-Antp) VS. Gb-abd-A Total | 115.0 | -3.0722 | 0.002 | Yes |
| (Gb-abd-A+ Gb-Antp) VS. Gb-Antp A1 | 523.0 | -0.68685 | 0.490 | No |
| (Gb-abd-A+ Gb-Antp) VS. Gb-Antp A2 | 366.0 | -2.41123 | 0.016 | Yes |
| (Gb-abd-A+ Gb-Antp) VS. Gb-Antp A3 | 305.0 | 3.17135 | 0.001 | Yes |
| (Gb-abd-A+ Gb-Antp) VS. Gb-Antp A4 | 412.0 | 1.82555 | 0.067 | No |
| (Gb-abd-A+ Gb-Antp) VS. Gb-Antp A5 | 483.5 | 0.94705 | 0.342 | No |
| (Gb-abd-A+ Gb-Antp) VS. Gb-Antp A6 | 532.0 | 0.34268 | 0.728 | No |
| (Gb-abd-A+ Gb-Antp) VS. Gb-Antp Total | 69.0 | 2.46699 | 0.013 | Yes |
| Gb-Antp VS. Gb-Ubx Total | 177.0 | -2.1157 | 0.034 | Yes |
| Gb-Antp VS. Gb-abd-A Total | 140.0 | -1.01006 | 0.312 | No |
| Gb-Ubx VS. Gb-abd-A Total | 461.0 | -0.67416 | 0.503 | No |

**Table S7:**
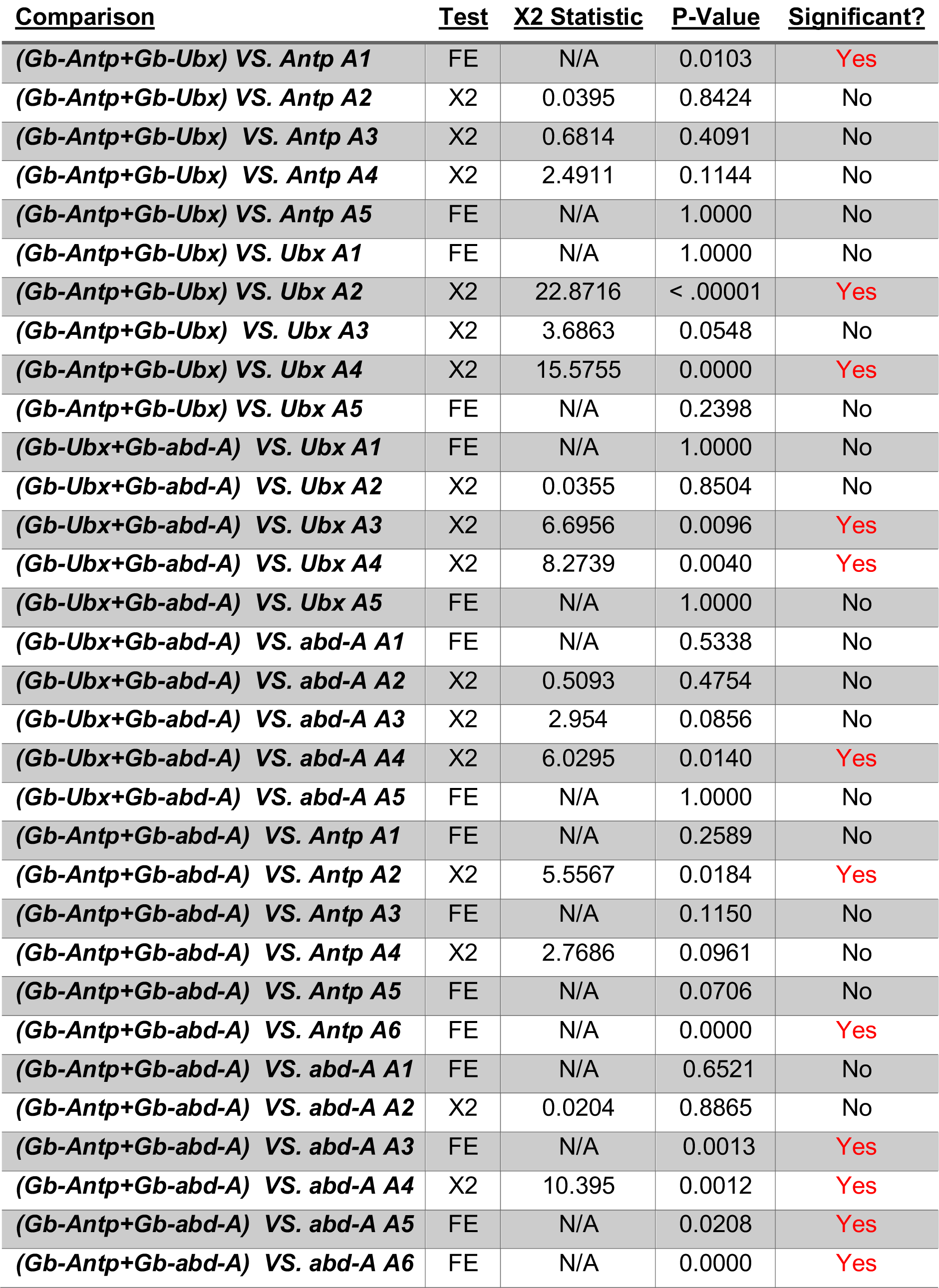
Statistical tests of presence/absence of PGC clusters in single vs. double Hox eRNAi treatments. FE = Fisher’s Exact Test; X2 = Chi-Square Test; NA = not applicable.

**Table S8:**
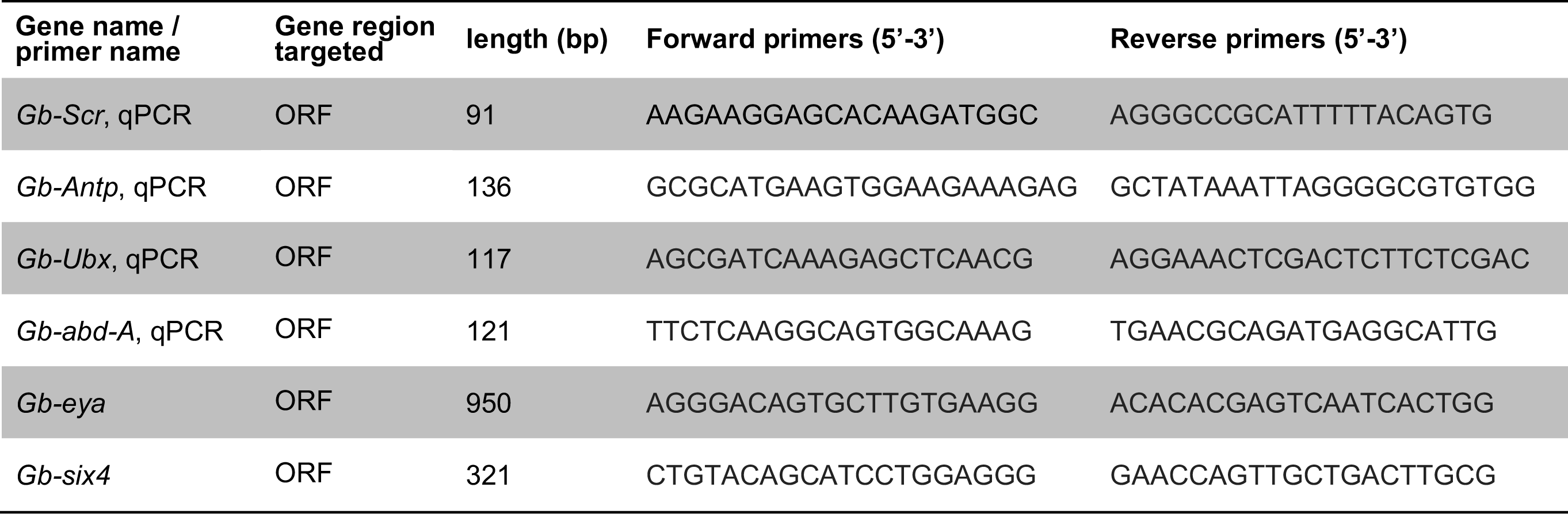
Primers used for gene cloning and qPCR.

